# MoChia1 is a family 18 glycoside hydrolase from the rice blast fungus, *Magnaporthe oryzae*, that releases reducing-end GlcNAc from chitin oligosaccharides

**DOI:** 10.64898/2025.12.09.693173

**Authors:** Takayuki Ohnuma, Syun Imaoka, Chikara Kataoka, Teruki Yoshimoto, Ryuta Okada, Toru Takeda, Tamo Fukamizo, Shohei Sakuda, Makoto Ogata

## Abstract

MoChia1, a family 18 glycoside hydrolase from the rice blast fungus *Magnaporthe oryzae*, was identified as an enzyme that binds chitin oligosaccharide elicitor to suppress the chitin-induced rice immune response. It hydrolyzed chitin oligosaccharides with a degree of polymerization of at least 3 [(GlcNAc)*_n_* (*n*≥3)] in an exo-splitting manner with anomeric retention. HPLC analysis of (GlcNAc)*_n_* cleavage by MoChia1 revealed that it specifically recognizes the *β*-anomer of the substrate at the + 1 subsite and cleaves the terminal glycosidic linkage at the reducing end. It also hydrolyzed the partially deacetylated chitotrisaccharide GlcN-(GlcNAc)_2_ to GlcN-GlcNAc and GlcNAc, but not (GlcN)_2_-GlcNAc, indicating the requirement of the *N*-acetyl group at the −1 subsite for the hydrolytic reaction. MoChia1 released *p*-nitrophenol from *p*NP-(GlcNAc)_2_, *p*-nitrophenyl di-*N*-acetyl-*β*-chitotrioside. Furthermore, it hydrolyzed the chitotriose derivatives with a modified GlcNAc residue on the reducing end, (GlcNAc)_2_-G, 4-*O*-*β*-di-*N*-acetylchitobiosyl-2-acetamido-2-deoxy-2,3-anhydro-glucopyranose and (GlcNAc)_2_-L, 4-*O*-*β*-di-*N*-acetylchitobiosyl-2-acetamido-2-deoxy-2,3-anhydro-glucono-δ-lactone, to (GlcNAc)_2_ and the modified GlcNAc, respectively. However, it did not hydrolyze 4-*O*-*β*-di-*N*-acetylchitobiosyl moranoline [(GlcNAc)_2_-M], reduced chitotriose (chitotriitol), and *α*-(GlcNAc)_2_ fluoride [(*α*-(GlcNAc)_2_-F]. MoChia1 did not bind to chitin and barely hydrolyzed the polymeric substrate, glycol chitin. Taken together, we concluded that MoChia1 is a GH18 reducing end GlcNAc-releasing exo-oligochitinase with the *β*-anomer selectivity. Allosamidin, a potent inhibitor of GH18 chitinases, was found to bind to MoChia1 and inhibit its hydrolytic activity with an *IC*_50_ of 54.4 ± 6.91 μM, indicating that allosamidin may be a potential candidate for a pesticide to prevent rice blast infection by inhibiting the chitinase activity of MoChia1.

## Introduction

Chitin, a linear polysaccharide comprised of *N*-acetyl-D-glucosamine units linked with *β*-1,4 linkages, is a major component of fungal cell walls. Together with other components, such as glucans and glycoproteins, it provides strength and shape to fungal cells (1–3). To break down the polysaccharides in their cell walls, fungi produce various glycoside hydrolases, including chitinolytic enzymes (4–6). These include chitinases, chitosanases, *β*-*N*-acetylglucosaminidases, chitin deacetylases, and lytic polysaccharide monooxygenases. Chitinase (EC 3.2.1.14) is a member of glycoside hydrolase families 18 and 19 (GH18 and GH19) according to the CAZy classification (www.cazy.org/) (7). It breaks the *β*-1,4 glycosidic linkages in chitin, converting them into shorter oligosaccharides. Fungi are well-known producers of GH18 chitinases but not GH19s (8,9). These enzymes are involved in several processes, including morphogenesis, cell division, autolysis, and the acquisition of chitin for nutritional purposes. Recently, it has been reported that GH18 chitinases, which are secreted by phytopathogenic fungi, play a significant role in the infection process in addition to their physiological functions (10,11). During the infection process of phytopathogens, chitin oligosaccharides with a longer chain length derived from their cell walls act as elicitors (12,13). These oligosaccharides are likely produced through the action of plant chitinases. Eventually, they are recognized by plant receptors called pattern-recognition receptors (PRRs), which are located on the plasma membrane, to transduce signals to downstream components and initiate immune responses (14,15). Therefore, GH18 enzymes from phytopathogens are believed to break them down into small fragments, rendering them unrecognizable to PRRs.

MoChia1, a GH18 chitinase produced by the rice blast fungus *Magnaporthe oryzae*, has been shown to function as a virulence factor (10). It was suggested to bind to chitin oligosaccharide elicitors and hide them from PRRs, thereby suppressing the chitin-triggered rice immune response during *M. oryzae* infection. Functions similar to those described have been reported in UvGH18.1, a GH18 chitinase from the fungus *Ustilaginoidea virens*, which is a phytopathogen that causes rice false smut. UvGH18.1 not only hydrolyzes chitin oligosaccharide elicitors, but also interacts with OsCEBiP, a rice chitin elicitor binding protein and the co-receptor OsCERK1, a rice chitin elicitor receptor kinase 1 (11). This impairs their chitin-induced dimerization and suppresses pathogen-associated molecular pattern (PAMP)-triggered immunity. Therefore, certain GH18 chitinases secreted by phytopathogens clearly play a significant role as a virulence factor in their infection process. However, their enzymatic properties have not yet been fully elucidated. In this study, we expressed, purified, and characterized the MoChia1 protein from *M. oryzae* to understand its role in evading the plant immune system in rice.

## Results

### Enzyme preparation and enzyme activity of MoChia1

MoChia1 was identified as a protein that *M. oryzae* secretes into the rice apoplast during infection. Analysis of the amino acid sequence of MoChia1 using the program SignalP predicted that it contains a signal sequence consisting of 26 amino acid residues at the N-terminus (16). Thus, an expression system for the predicted mature protein consisting of 371 amino acid residues (27–397) fused to an N-terminal 6 × His tag was created and the recombinant MoChia1 protein was produced in *Escherichia coli*. The yield of recombinant MoChia1 was approximately 30 mg from 1 liter of induced culture. From the SDS-PAGE analysis, purified MoChia1 had a molecular mass of 43 kDa, which corresponded with the molecular mass calculated from the amino acid sequence (*M*_r_ = 42132.27) (Fig. S1). MoChia1 hydrolyzed *p*NP-(GlcNAc)_2_ and (GlcNAc)*_n_* (*n* = 3–6), but not (GlcNAc)_2_. The enzymatic properties were determined as a function of pH and temperature using *p*NP-(GlcNAc)_2_ (Fig. 1). The optimum pH for chitinase activity was approximately 5.0 and the optimum temperature was 40 °C (Figs. 1A and 1B). MoChia1 was stable between pH 5.0 and 9.5, but unstable below pH 5.0 and above pH 9.5. With respect to temperature, MoChia1 retained 80% of its activity up to 40°C and became unstable above 40°C. Among the natural glycosides tested, MoChia1 produced more reducing sugars as the oligosaccharide chain length increased (Table 1). However, it hardly hydrolyzed glycol chitin, an ethylene glycol derivative of chitin in which *O*-6 of the chitin chain was etherified (data not shown).

**Figure 1.**
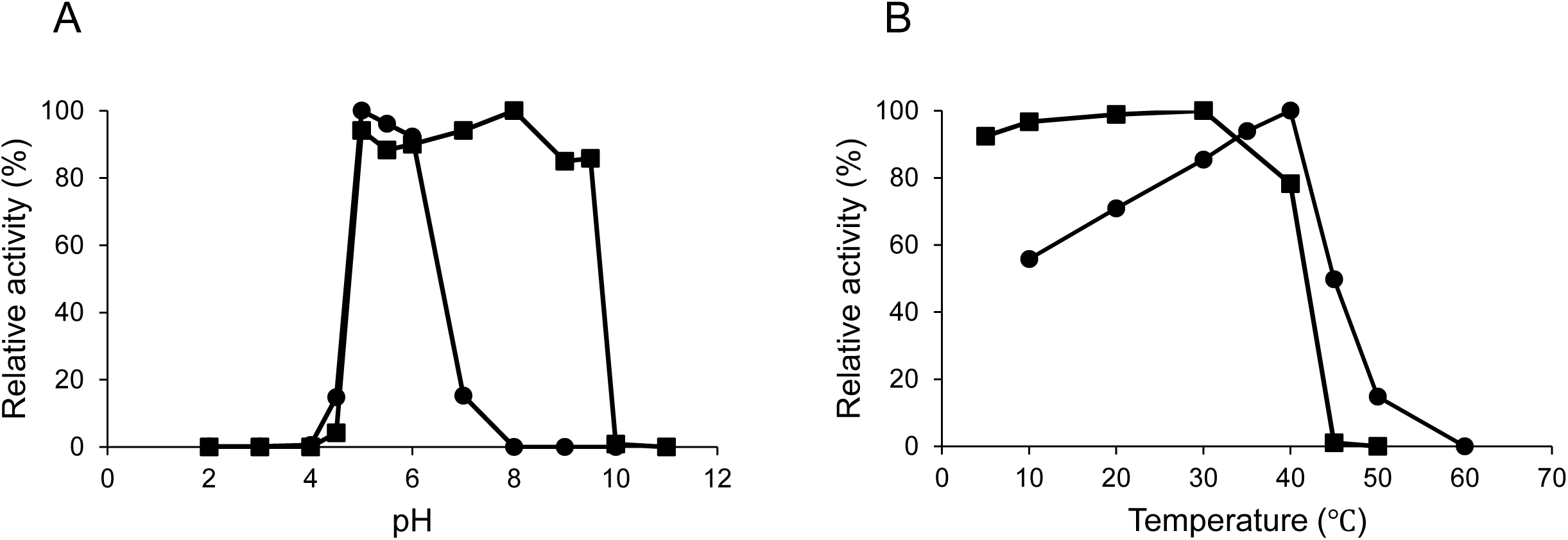
Effect of pH and temperature on activity and stability of MoChia1. (*A*), effect of pH on MoChia1 activity (closed circles) and pH stability (closed squares) was determined using various 20 mM buffers (glycine-HCl buffer pH, 2.0–3.0; sodium-acetate buffer pH, 3.5–5.5; sodium-phosphate buffer pH, 6.0–7.5; Tris-HCl, 8.0–9.0; glycine-NaOH buffer, 9.5–10.5; KCl-NaOH buffer, 11–12). For pH stability, MoChia1 was preincubated in various pH buffers at 37°C for 24 h. The residual activity was assessed under standard assay conditions. (*B*), effect of temperature on MoChia1 activity (closed circles) and thermostability (closed squares) was determined at various temperatures (5–60°C) at pH 5.0. For thermostability, MoChia1 was preincubated at various temperatures for 0.5 h. The residual activity was assessed under standard assay conditions. *p*NP-(GlcNAc)_2_ was used as a substrate. Liberated *p*NP was quantified spectrophotometrically at 405 nm.

**Table 1.** Specific activity of MoChia1.

| Substrate | Specific activity (U/mg)* | Relative activity (%) |
| --- | --- | --- |
| (GlcNAc)<sub>2</sub> | N.D. | N.D. |
| (GlcNAc)<sub>3</sub> | 0.59 | 20 |
| (GlcNAc)<sub>4</sub> | 1.57 | 54 |
| (GlcNAc)<sub>5</sub> | 2.30 | 79 |
| (GlcNAc)<sub>6</sub> | 2.92 | 100 |

\*Specific activity was defined as $\mu\text{mol}/\text{min}$ of reducing sugar equivalent to GlcNAc released per milligram of enzyme.

### TLC analysis of the reaction products produced by MoChia1 from (GlcNAc)_n_

We analyzed the time-dependent degradation of (GlcNAc)*_n_* (*n* = 2–6) by MoChia1 using TLC (Fig. 2A–2E). The TLC analysis clearly indicated that MoChia1 acted exolytically, yielding GlcNAc and (GlcNAc)_2_ as the end products from chitin oligosaccharides with a degree of polymerization of at least 3 [(GlcNAc)*_n_* (*n*≥3)]. (GlcNAc)*_n_* (*n* = 3–6) was hydrolyzed to (GlcNAc)*_n_*_-1_ + GlcNAc and the intermediates (GlcNAc)*_n_*_-1_, which were one GlcNAc unit shorter than the starting substrates, were further hydrolyzed to the end products GlcNAc and (GlcNAc)_2_. We also investigated the hydrolytic activity of MoChia1 with respect to the partially deacetylated chitotrisaccharide GlcN-(GlcNAc)_2_ and (GlcN)_2_-GlcNAc. In addition to (GlcNAc)*_n_*, these heterotrisaccharides may be released from the cell wall of *M. oryzae*, making them potential substrates for MoChia1 in nature (17,18). MoChia1 hydrolyzed GlcN-(GlcNAc)_2_ to GlcN-GlcNAc and GlcNAc, but not (GlcN)_2_-GlcNAc, suggesting that the *N*-acetyl group at the –1 subsite is required for the hydrolytic reaction, but not at the –2 subsite. (Figs. 3A and 3B).

**Figure 2.**
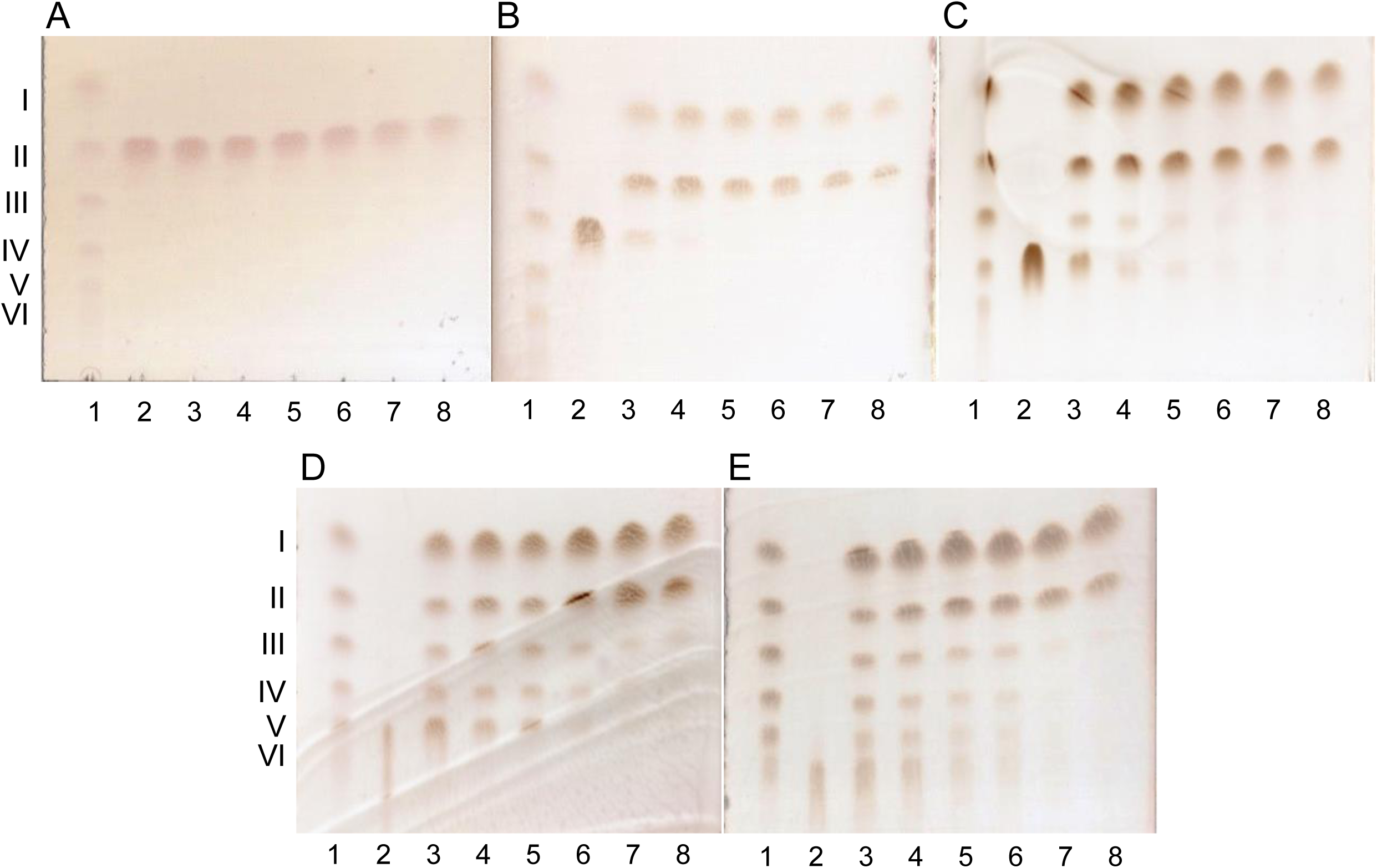
TLC analysis of (GlcNAc)*_n_* (*n* = 2–6) hydrolysis by MoChia1. A reaction mixture comprising 2.0 μM MoChia1 and 8.0 mM (GlcNAc)_2_ (*A*), (GlcNAc)_3_ (*B*), (GlcNAc)_4_ (*C*), (GlcNAc)_5_ (*D*), and (GlcNAc)_6_ (*E*) in 20 mM sodium acetate buffer (pH 5.0) was incubated at various times (from 0.5 to 3 h) at 37 °C. (*A*–*E*), Lanes: 1, I–VI represent a standard mixture of (GlcNAc)*_n_* (*n* = 1–6); 2, substrate; 3–8, incubation for 0.5, 1.0, 1.5, 2.0, 2.5, and 3.0 h, respectively.

**Figure 3.**
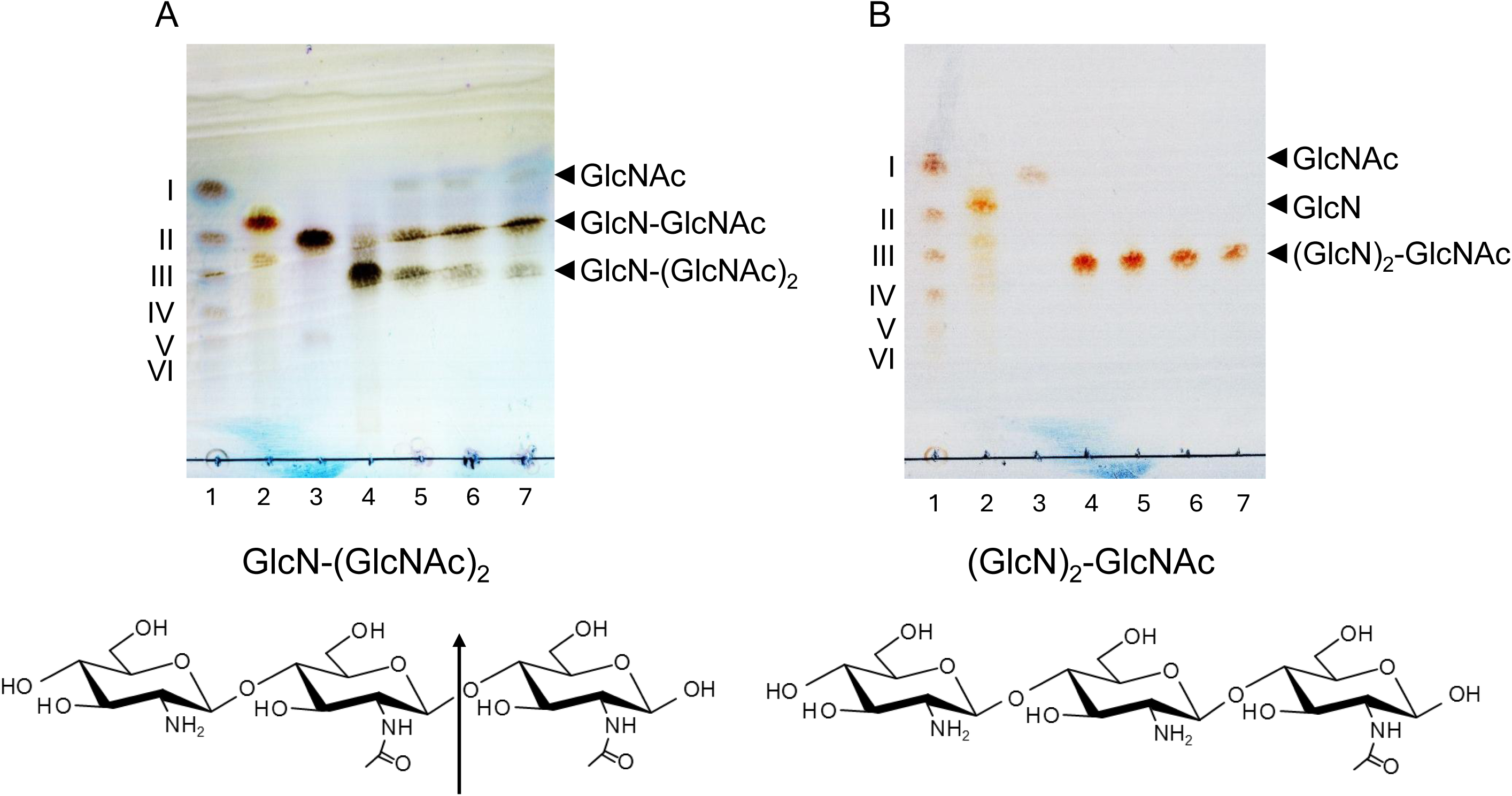
TLC analysis of the partially deacetylated chitotrisaccharide hydrolysis by MoChia1. A reaction mixture comprising 2.0 μM MoChia1 and 8.0 mM GlcN-(GcNAc)_2_ (*A*) and (GlcN)_2_-GlcNAc (*B*) in 20 mM sodium acetate buffer (pH 5.0) was incubated at various times (from 1.0 to 3.0 h) at 37°C. (*A*), Lanes: 1, I–VI represent a standard mixture of (GlcNAc)*_n_* (*n* = 1–6); 2, GlcN; 3, GlcN-GlcNAc; 4, substrate; 5–7, incubation for 1.0, 2.0, and 3.0 h, respectively. (*B*), Lanes: 1, I–VI represent a standard mixture of (GlcNAc)*_n_* (*n* = 1–6); 2, GlcN; 3, GlcNAc; 4, substrate; 5–7, incubation for 1.0, 2.0, and 3.0 h, respectively. The chemical structure of each substrate is shown below each TLC plate.

### Anomer analysis of the reaction products produced by MoChia1 from (GlcNAc)_n_

The anomeric form of the enzymatic reaction product provides information about the cleavage site of the oligosaccharide substrate. HPLC profiles depicting the time-dependent enzymatic hydrolysis of (GlcNAc)_3_ and (GlcNAc)_5_ are shown in Fig. 4. The initial products of substrate (GlcNAc)_3_ hydrolysis were GlcNAc and (GlcNAc)_2_, which were the *β-*anomers, although they gradually mutarotated to *α-*anomers. Furthermore, the HPLC profile of this reaction showed that the peak area of the *β*-anomer of (GlcNAc)_3_ decreased over time, whereas the area of the *α*-anomer remained almost unchanged (Fig. 4A). In the case of (GlcNAc)_5_ hydrolysis, the initial products were exclusively GlcNAc and (GlcNAc)_4_, which were the *β-*anomers. The *β*-anomer of (GlcNAc)_5_ decreased apparently as the reaction proceeded (Fig. 4B). The product (GlcNAc)_4_ generated from (GlcNAc)_5_ was further hydrolyzed to *β*-anomers of GlcNAc and (GlcNAc)_3_ as the substrate (GlcNAc)_4_ was hydrolyzed (Fig. S2A). The (GlcNAc)_6_ substrate was hydrolyzed, yielding *β*-anomer GlcNAc and (GlcNAc)₅, which were subsequently degraded in the same manner (Fig. S2B). Because the enzymatic reaction takes place through a substrate-assisted catalysis mechanism in GH18 family enzymes, the anomeric form of the newly formed reducing end residue obtained from MoChia1-catalyzed hydrolysis of (GlcNAc)*_n_* is always the *β*-anomer. Therefore, these results indicate that MoChia1 specifically recognizes the *β*-anomer of the chitin oligosaccharide substrate at the +1 subsite and hydrolyzes the first glycosidic linkage from the reducing end of the substrate.

**Figure 4.**
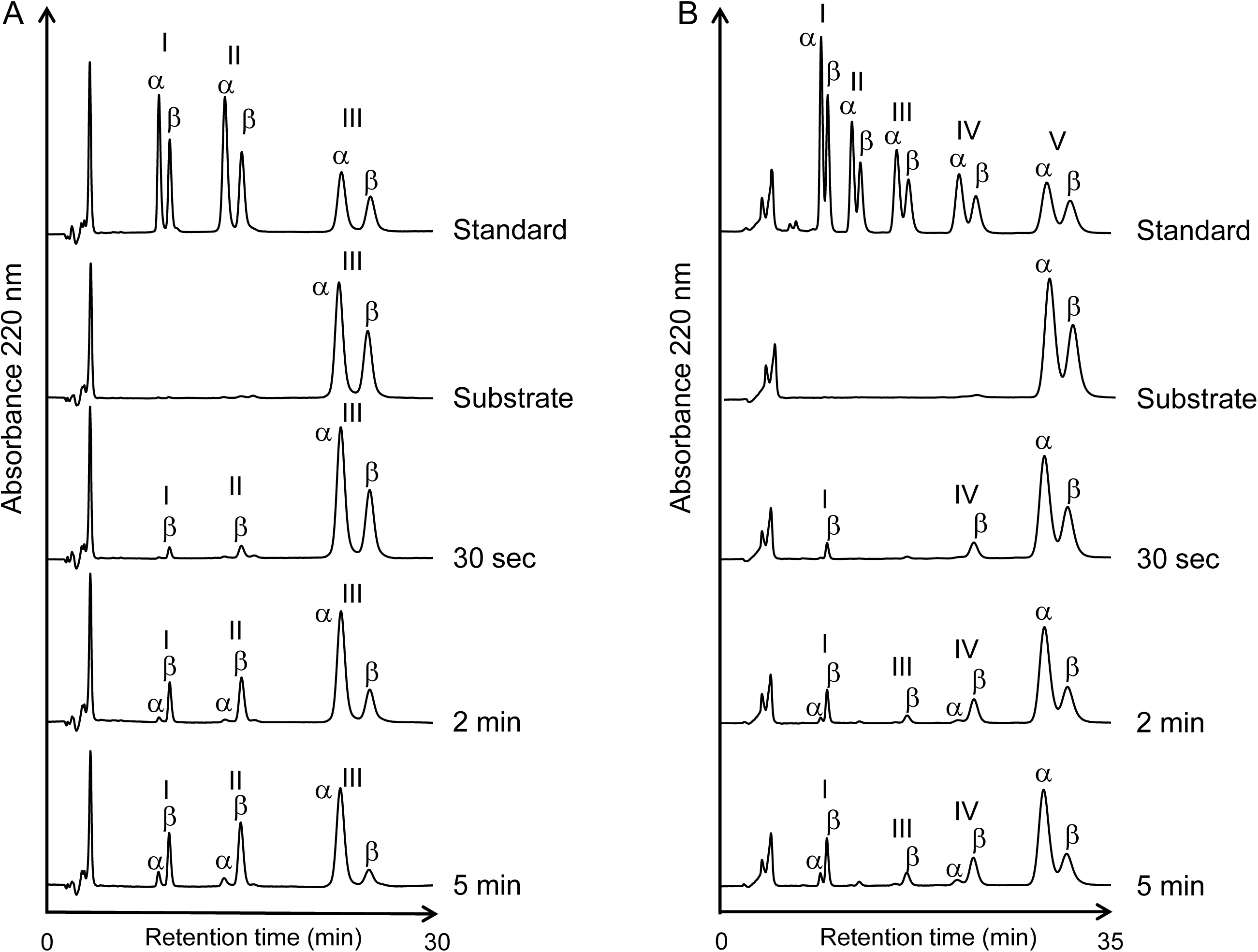
Anomer formation catalyzed by MoChia1 from the (GlcNAc)_3_ and (GlcNAc)_5_ substrates. Time-dependent HPLC profiles showing the hydrolysis of (GlcNAc)_3_ (*A*) and (GlcNAc)_6_ (*B*) by MoChia1. Enzyme and substrate concentrations were 2.0 μM and 8 mM, respectively. The enzyme reaction was conducted in 20 mM sodium acetate buffer (pH 5.0) at 25°C. Numerals in the figure represent the degree of polymerization.

### Substrate specificity

The substrate specificity of MoChia1 was investigated by examining the hydrolytic activity of the enzyme using various (GlcNAc)_2_ and (GlcNAc)_3_ derivatives. When 4-*O*-*β*-di-*N*-acetylchitobiosyl-2-acetamido-2-deoxy-2,3-anhydro-glucopyranose [(GlcNAc)_2_-G] was used as the substrate, it was hydrolyzed into (GlcNAc)_2_ and 2-acetamido-2-deoxy-2,3-anhydro-glucopyranose (G) (Fig. 5A). In the case of 4-*O*-*β*-di-*N*-acetylchitobiosyl-2-acetamido-2-deoxy-2,3-anhydro-glucono-δ-lactone [(GlcNAc)_2_-L], it was hydrolyzed into (GlcNAc)_2_ and 2-acetamido-2-deoxy-2,3-anhydro-glucono-δ-lactone (L). A portion of L was spontaneously converted to unidentified compound X (Fig. 5B). *p*NP-(GlcNAc)_2_ was hydrolyzed into (GlcNAc)_2_ and *p*-nitrophenol (Figs. 5C and 5D). However, 4-*O*-*β*-di-*N*-acetylchitobiosyl moranoline [(GlcNAc)_2_-M], reduced chitotriose (chitotriitol), and *α*-(GlcNAc)_2_ fluoride [(*α*-(GlcNAc)_2_-F] were not hydrolyzed (Fig. S3). These results also revealed that MoChia1 hydrolyzes the reducing end glycosidic linkage of oligosaccharide substrates. Recognition of substrates at the +1 subsite of MoChia1 has been shown to be less stringent, as evidenced by the recognition of G, L, and *p*-nitrophenol residues at this site.

**Figure 5.**
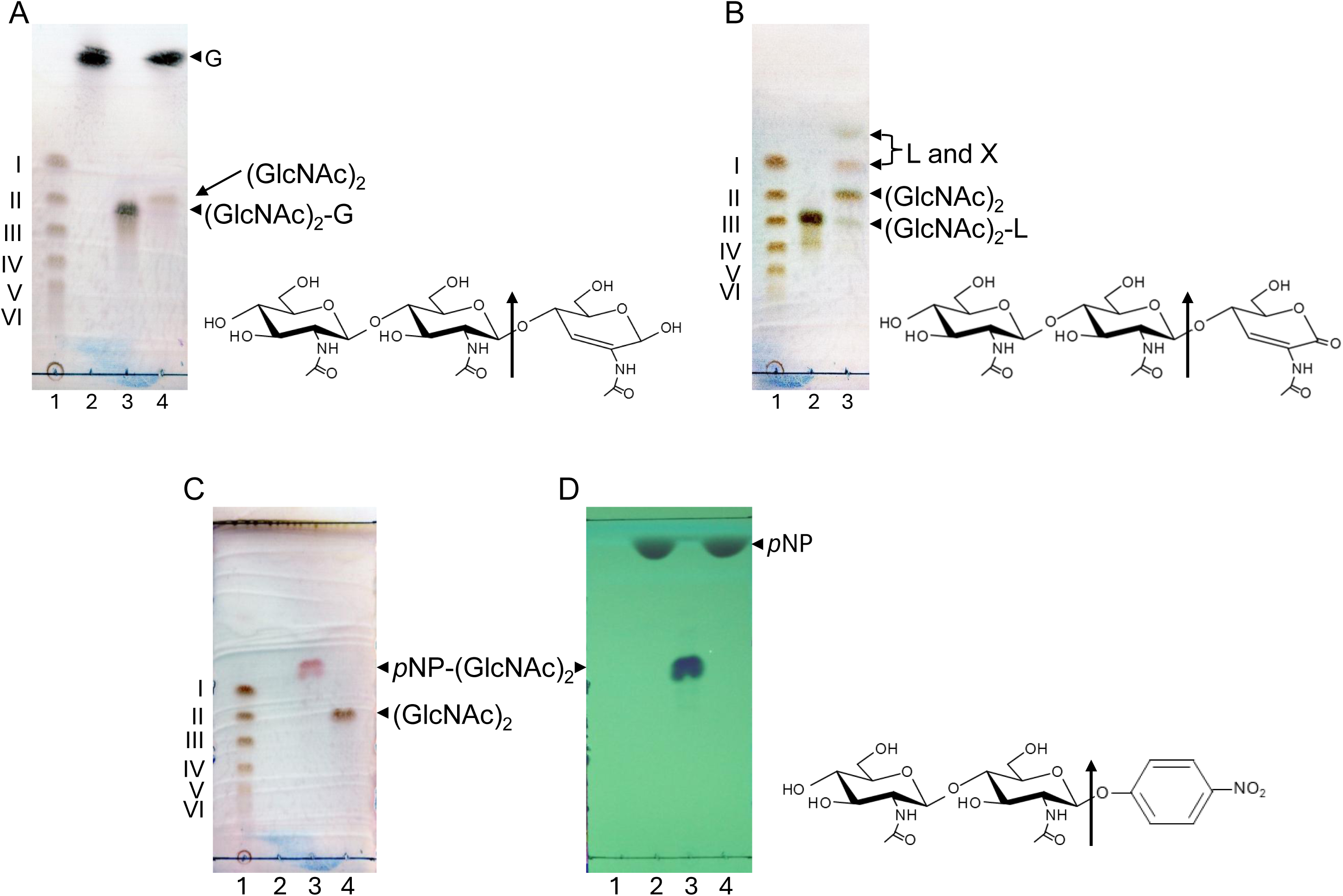
TLC analysis of the hydrolysis of the chitotriose derivatives modified with a modified GlcNAc residue on the reducing end and *p*NP-(GlcNAc)_2_ by MoChia1. A reaction mixture comprising 2.0 μM MoChia1 and 8.0 mM (GlcNAc)_2_-G, (GlcNAC)_2_-L, and *p*NP-(GlcNAc)_2_ in 20 mM sodium acetate buffer (pH 5.0) was incubated for 60 min at 37°C. Reaction products were detected with methanol containing sulfuric acid (*A*–*C*) and visualized under UV light at 254 nm (*D*). (*A*), Lanes: 1, I–VI represent a standard mixture of (GlcNAc)*_n_* (*n* = 1–6); 2, G (2-acetamido-2-deoxy-2,3-anhydro-glucopyranose); 3, (GlcNAc)_2_-G; 4, reaction product. (*B*), Lanes: 1, I–VI represent a standard mixture of (GlcNAc)*_n_* (*n* = 1–6); 2, (GlcNAc)_2_-L; 3, reaction product. (*C*, *D*), Lanes: 1, I–VI represent a standard mixture of (GlcNAc)*_n_* (*n* = 1–6); 2, *p*-nitrophenol; 3, *p*NP-(GlcNAc)_2_; 4, reaction product. The chemical structure of each substrate is shown to the right of each TLC plate.

### Kinetic characterization of MoChia1 and inhibitory effect of allosamidin

Since (GlcNAc)_2_-G was only hydrolyzed once by MoChia1 into (GlcNAc)_2_ and G, and it is more soluble than *p*NP-(GlcNAc)_2_, reaction kinetics were analyzed using this synthetic substrate. The initial velocity profiles of G release catalyzed by MoChia1 at different (GlcNAc)_2_-G concentrations (0–20 mM) are shown in Fig. 6A. The kinetic parameters were calculated as listed in Table 2. MoChia1 was assigned a *K*_m_ value of 9.74 ± 2.34 mM and a *k*_cat_ value of 5.0 ± 0.53 s^−1^. The catalytic efficiency, *k*_cat_/*K*_m_, was 0.51 ± 0.13 s^−1^ mM^−1^. To determine the inhibitory effect of allosamidin on MoChia1, the percentage inhibitions were plotted against the concentrations of allosamidin. As shown in Fig. 6B, allosamidin inhibited MoChia1 activity in a dose-dependent manner, with an *IC*_50_ value of 54.4 ± 6.91 μM (Table 2).

**Figure 6.**
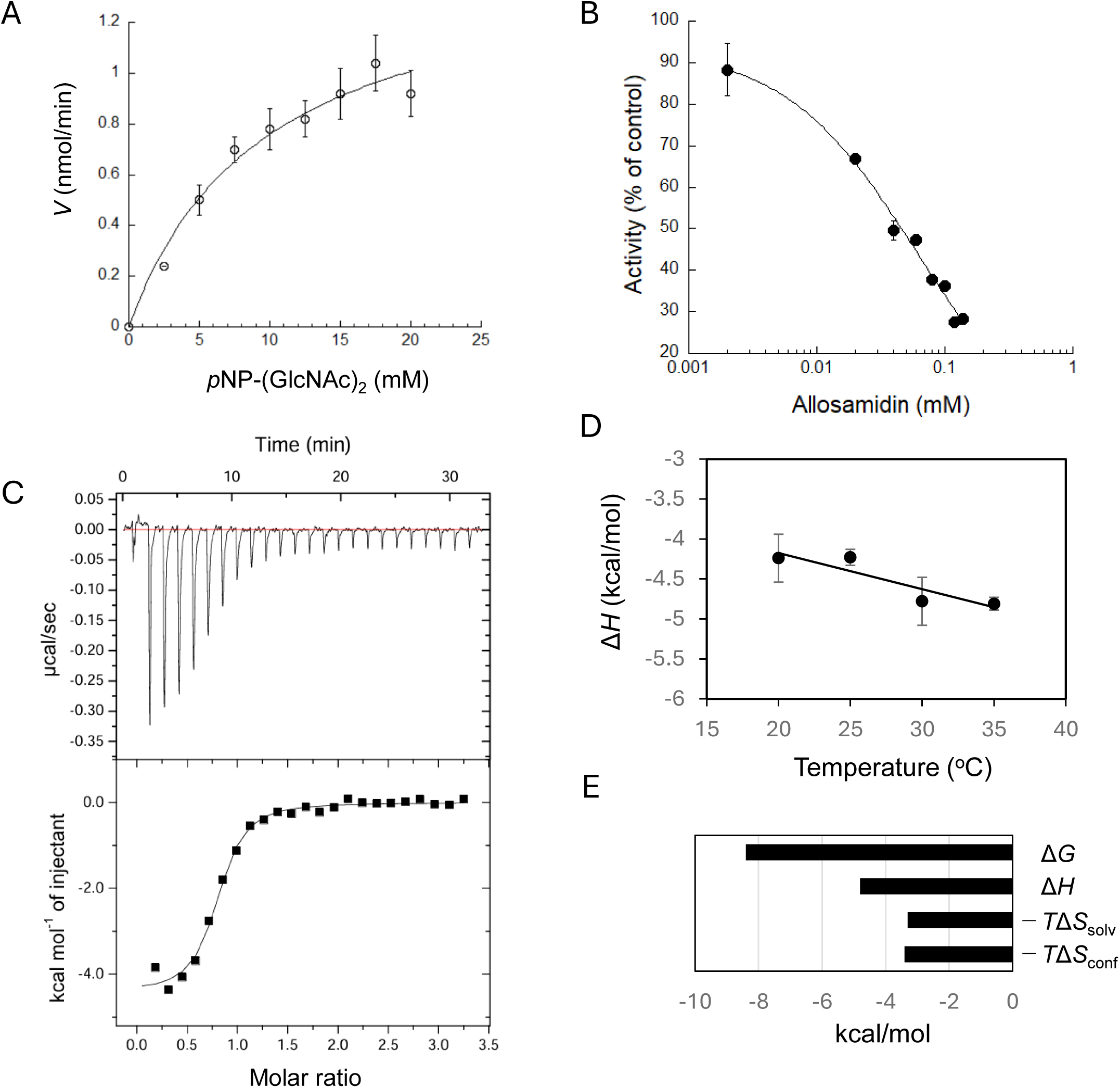
Analysis of the inhibition of MoChia1 by allosamidin. (*A*), the Michaelis–Menten plot shows the kinetics of MoChia1 with *p*NP-(GlcNAc)_2_ as substrate. (*B*), dose-response curve of MoChia1 inhibition for allosamidin. (*C*), binding isotherms (top) with theoretical fits (bottom) for binding of allosamidin to MoChia1. (*D*), temperature dependence of the enthalpy change for binding of allosamidin to MoChia1. The plots of Δ*H*° versus temperature yielded a change in heat capacity (Δ*C*_p_°) based on the slope of the line. (*E*), the bar diagram of thermodynamic parameters for binding of allosamidin to MoChia1. Error bars represent standard deviations obtained from three independent experiments.

**Table 2.** Kinetic characterization of MoCHia1 and the inhibitory effect of allosamidin.

| Enzyme | $K_m$ (mM) | $k_{\text{cat}}$ ($\text{s}^{-1}$) | $k_{\text{cat}}/K_m$ ($\text{s}^{-1}\text{mM}^{-1}$) | $IC_{50}$ ($\mu\text{M}$) |
| --- | --- | --- | --- | --- |
| MoChia1 | $9.74 \pm 2.34$ | $5.0 \pm 0.53$ | $0.51 \pm 0.13$ | $54.4 \pm 6.91$ |

### ITC analysis of allosamidin binding to MoChia1

The interaction between allosamidin and MoChia1 was studied using ITC at 30°C and pH 7.0. Fig. 6C shows an ITC thermogram and the theoretical fit to the experimental data for allosamidin. The theoretical fit was successfully obtained based on the experimental data using a nonlinear least-squares algorithm by varying the binding affinity constant (*K*_a_), number of binding sites (i.e., stoichiometry of the reaction (*N*)), and enthalpy change of ligand binding (Δ*H*°). The titration curve fit well to a single binding model, with a calculated *N* of 0.76 ± 0.07, indicating a binding stoichiometry of 1:1. At this temperature and pH, allosamidin was found to bind to MoChia1 with a *K*_d_ of 0.98 ± 0.22 μM. Binding was confirmed to be both enthalpically (Δ*H*° = −4.8 ± 0.3 kcal/mol) and entropically (−*T*Δ*S*° = −3.6 ± 0.4 kcal/mol) driven, resulting in a Δ*G*° of −8.4 ± 0.1 kcal/mol (Table 3).

**Table 3.** Thermodynamic parameters for the binding of allosamidin to MoChia1 at 30 °C.

| $N$ | $K_d$ ($\mu\text{M}$) | $\Delta H$ (kcal/mol) | $-T\Delta S$<br>(kcal/mol) | $\Delta G$ (kcal/mol) |
| --- | --- | --- | --- | --- |
| $0.76 \pm 0.07$ | $0.98 \pm 0.22$ | $-4.8 \pm 0.3$ | $-3.6 \pm 0.4$ | $-8.4 \pm 0.1$ |

### Heat capacity change and parameterization of the entropic term

The parameterization of the entropy change upon allosamidin binding to MoChia1 provides insight into the binding mechanism. The Δ*C*_p_° value for the binding interaction was calculated using ITC titrations over a temperature range from 20°C to 35°C. The temperature dependence of Δ*H*° for binding in this temperature range yielded a straight line with a slope of −45.2 ± 15.3 cal/K·mol (Δ*C*_p_°) at pH 7.0 (Fig. 6D and Table 4), resulting the favorable solvation entropy change for binding (−*T*Δ*S*_solv_° = −3.3 ± 1.1 kcal/mol). The contribution of conformational entropy change to binding was virtually the same extent (−*T*Δ*S*_conf_° = −3.4 ± 1.1 kcal/mol) (Fig. 6E and Table 4).

**Table 4.** Heat capacity change (Δ*C*_p_°) for the binding of allosamidin to MoChia1 and dissection of the entropic term.

| $\Delta C_p$ (cal/K mol) | $-T\Delta S$<br>(kcal/mol) | $-T\Delta S_{\text{solv}}$<br>(kcal/mol) | $-T\Delta S_{\text{mix}}$<br>(kcal/mol) | $-T\Delta S_{\text{conf}}$<br>(kcal/mol) |
| --- | --- | --- | --- | --- |
| $-45.2 \pm 15.3$ | $-4.3 \pm 0.2$ | $-3.3 \pm 1.1$ | 2.4 | $-3.4 \pm 1.1$ |

### Chitin binding assay

We used pull-down binding assays to examine whether MoChia1 directly binds to insoluble chitin. As shown in Fig. 7, *Sm*ChiB, a protein from *Serratia marcescens* consisting of a chitin-binding domain and a GH18 catalytic domain, bound to chitin. It was then precipitated with chitin by centrifugation. Most of the input protein disappeared from the liquid fraction. On the other hand, MoChia1 remained in the supernatant after centrifugation and can barely be detected in the bound fraction as BSA. Based on these results, we concluded that MoChia1 is unable to bind to chitin.

**Figure 7.**
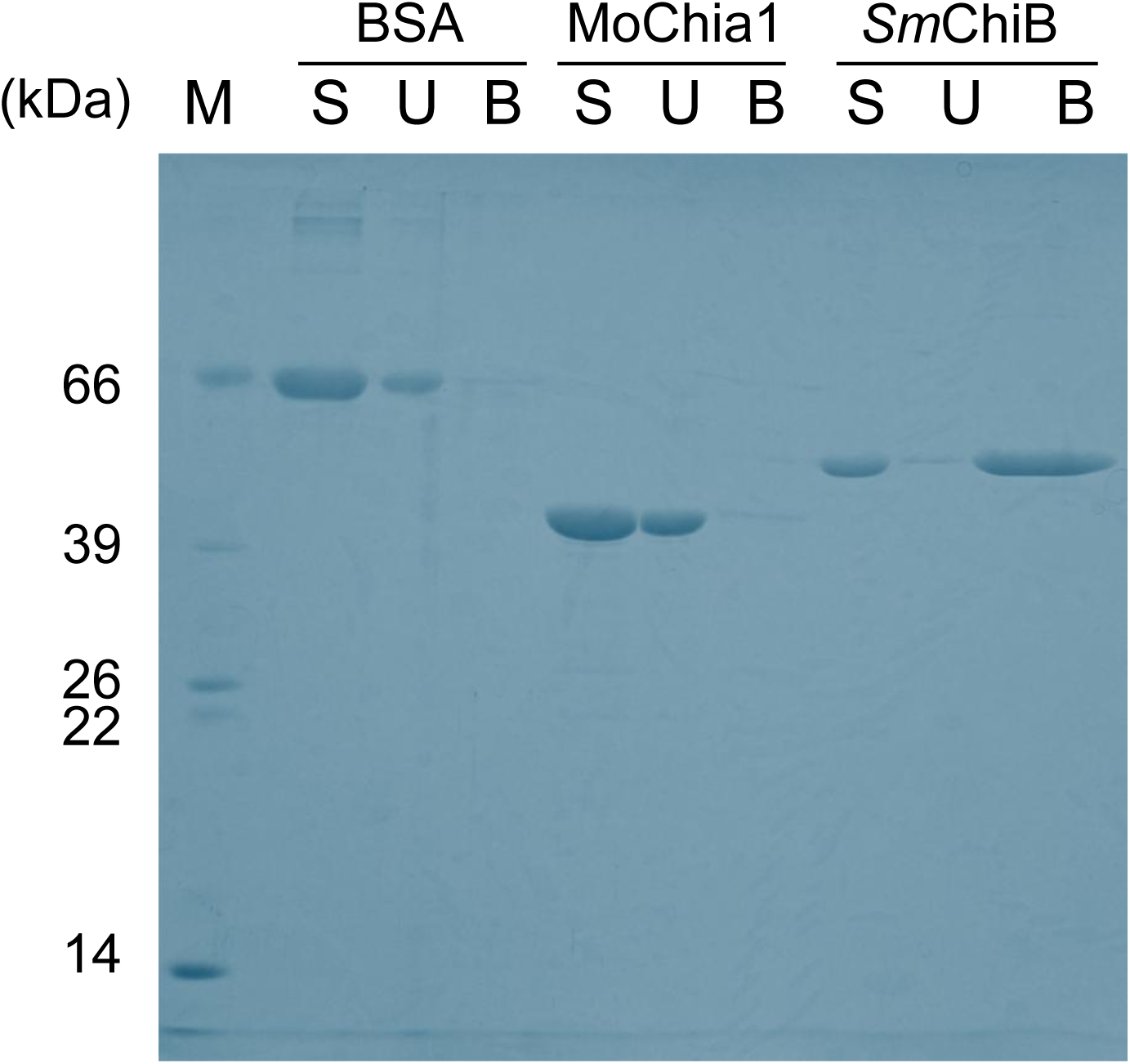
Chitin binding assay. Chitin binding activity of MoChia1 was assessed by incubating the protein with chitin beads. The fractions were then analyzed on SDS-PAGE. *Sm*ChiB was used as a positive control and BSA as a negative control. S, start fraction representing the amount of protein loaded onto the chitin beads; U, unbound protein (supernatant fraction); B, bound protein (pellet fraction).

## Discussion

MoChia1 was isolated from the growth culture of *M. oryzae* during the screening process for new elicitors/PAMPs that trigger the rice immune response. It has been demonstrated that MoChia1 binds to the chitin oligosaccharide elicitor during the infection process. This suppresses the chitin-induced reactive oxygen species burst in rice cells and is thought to inhibit the recognition of the oligosaccharide elicitor by OsCEBIP and OsCERK1 (10). However, since MoChia1 is an active chitinase, we expected its hydrolytic activity would be involved in some way in the infection of *M. oryzae*. By using chitin oligosaccharides and their derivatives bearing various modifications on the reducing end GlcNAc residue as substrates, we examined the mode of action and specificity of MoChia1. It was revealed that MoChia1 does hydrolyze the linkage at the reducing end of the chitin oligosaccharide substrate with the *β*-anomer selectivity, releasing GlcNAc from that end. However, it does not hydrolyze the linkage at the non-reducing end, unlike GH20 *β*-*N*-acetylglucosaminidase (EC 3.2.1.52). It also differs from both endo-chitinase (EC 3.2.1.14) and exo-chitinase (EC 3.2.1.201). The latter was assigned as the enzyme that hydrolyzes the second glycosidic linkage from the reducing end of chitin, releasing (GlcNAc)_2_ in Enzyme Nomenclature (https://iubmb.qmul.ac.uk/enzyme/EC3/2/1/201.html). MoChia1 barely hydrolyzed glycol chitin and failed to bind to insoluble chitin. Considering these, we propose that MoChia1 is a novel GH18 chitinase and that it should be named reducing end GlcNAc-releasing exo-oligochinase (Rec). This name is reminiscent of Rex, a GH8 reducing end xylose-releasing exo-oligoxylanase, found in *Bacillus halodurans* C-125 (19).

Di-*N*-acetylchitobiase (also known as chitobiase, [CTBS]) is a lysosomal GH18 exo-glycosidase that hydrolyzes the *β*-1,4 glycosidic linkage of the chitobiose core of *N*-linked oligosaccharide (20). CTBS requires prior cleavage of the adjacent Asn-GlcNAc amide bond in order to act on it. This enzyme is the first report of GH18 chitinase that acts on the glycosidic linkage at the reducing end of the substrate, releasing GlcNAc. Later on, CTBS was reported to hydrolyze chitin oligosaccharides with a degree of polymerization of at least 2 [(GlcNAc)*_n_* (*n* ≧ 2)] from their reducing end (21,22). Notably, it was revealed that CTBS only recognizes the *α*-anomer of (GlcNAc)*_n_* (*n*≥2), releasing GlcNAc without hydrolyzing the *β*-anomer. A synthetic *p*-nitrophenyl glycoside, *p*NP-GlcNAc, was also not degraded by this enzyme. We do not yet know whether MoChia1 hydrolyzes the *β*-1,4 glycosidic linkage of the chitobiose core of *N*-linked oligosaccharides, though it does hydrolyze the *β*-anomer of (GlcNAc)*_n_* (*n*≥3)] and *p*NP-(GlcNAc)_2_. Because the anomeric configurations of the substrate and product are the same in the *β*-form, MoChia1 may hydrolyze the chitin oligosaccharide substrate processively. However, this is likely not the case for CTBS, since the configurations are opposite. Based on their amino acid sequences, these two proteins are predicted to have the same domain structure, consisting of a (*β*/*α*)_8_ fold with a small insertion domain composed of an *α*-helix and *β*-strands similar to the structures of other GH18 chitinases, but the identity between MoChia1 and CTBS is quite low (ca. 12%) (Fig. S4) (23). Therefore, it is very interesting to elucidate the substrate recognition mechanism of these two exo-GH18 chitinases, especially at the +1 subsite.

Van Munster et al. reported the recombinant production, purification and characterization of MBP-fused Cfcl, which is a GH18 chitinase from *Aspergillus niger*. They demonstrated that this enzyme cleaves the glycosidic linkage of (GlcNAc)_4_ at the reducing end, releasing GlcNAc (24). This is the second example of GH18 that acts on the glycosidic linkage at the reducing end. MBP-fused Cfcl hydrolyzed *p*NP-(GlcNAc)_2_ and *p*NP-(GlcNAc)_3_, releasing *p*-nitrophenol, but it did not hydrolyze (GlcNAc)_5_ alditol. Although MoChia1 hydrolyzed (GlcNAc)*_n_* (*n*≥3) and *p*NP-(GlcNAc)_2_ as CfcI, these two proteins have only 40% identical amino acids in their sequences. Of note, Cfcl has an unusual domain structure; that is, the carbohydrate-binding module 18 (CBM18) is integrated into the (*α*+*β*) insertion domain within the GH18 catalytic domain, which is absent in MoChia1 (Fig. S4). The *α*/*β*-anomer selectivity on chitin oligosaccharide substrate of Cfcl, as well as the substrate specificity at the +1 subsite, remains unknown.

Allosamidin inhibited the chitinase activity of MoChia1 with moderate inhibitory activity (*IC*_50_ = 54.4 ± 6.91 μM). The interaction between the allosamidin and GH18 enzymes of bacterial and plant origin has been extensively studied by using ITC (25,26). The driving force behind the interaction between *Sm*ChiA, a GH18 enzyme from *Serratia marcescens*, and allosamidin at 30°C was the significant enthalpic contribution (Δ*H*° = −7.8 to −6.2 kcal/mol) and small entropic contribution (−*T*Δ*S*° = −3.2 to −1.8 kcal/mol) to the Gibbs free energy (Δ*G*° = −9.7 to −9.4 kcal/mol). The main force between *Sm*ChiB, another type of GH18 enzyme found in *S. marcescens*, and allosamidin was the large entropic contribution (−*T*Δ*S*° = −13.2 to −9.2 kcal/mol) with an unfavorable or slightly favorable enthalpic contribution (Δ*H*° = −1.2 to 3.8 kcal/mol). Both enzymes have a catalytic domain for breaking down chitin and a separate chitin-binding domain help them bind to crystalline chitin substrates. In contrast, the plant GH18 chitinases (NtChiV from *Nicotiana tabacum*, AtChiC from *Arabidopsis thaliana*, and CrChiV from *Cycas revoluta*) have the same domain organization as MoChia1. The driving force behind the interaction between these plant GH18 enzymes and allosamidin at the same temperature was the combined enthalpic and entropic contributions (Δ*H*° = −6.5 to −4.5 kcal/mol and −*T*Δ*S*° = −5.1 to −2.9 kcal/mol). In these cases, the primary contributor to the interaction in the entropic term was the solvation entropy change (−*T*Δ*S*_solv_° = −9.8 to −4.8 kcal/mol), accompanied by an unfavorable or only slightly favorable conformational entropy change (−*T*Δ*S*_conf_° = −0.9 to 2.3 kcal/mol). The driving force for allosamidin binding to MoChia1 was the combined enthalpic and entropic contributions (Δ*H*° = −4.8 ± 0.3 kcal/mol and −*T*Δ*S*° = −3.6 ± 0.4 kcal/mol). However, in this case, both the solvation and conformational entropy changes contributed to the binding to a similar degree (−*T*Δ*S*_solv_° = −3.3 ± 1.1 kcal/mol and −*T*Δ*S*_conf_° = −3.4 ± 1.1 kcal/mol) (Fig. 6E and Table 4). A structural comparison of MoChia1 and CrChi-A revealed that MoChia1 has four extended loop structures (loop 1 to 4) (Figs. S5C and S5D). Furthermore, the wall structure formed by two of these loops (loops 2 and 3) blocks the aglycon binding site and creates part of the deep substrate binding pocket (Fig. S5B). The substrate binding cleft of CrChi-A is wide and open, exposing it to the solvent (Fig. S5A). Therefore, it is reasonable to assume that CrChi-A’s substrate binding site is more solvated than that of MoChia1 and repels more water molecules upon allosamidin binding. Furthermore, the binding of allosamidin to MoChia1 was more strongly influenced by changes in conformational entropy than by those of CrChi-A. Although allosamidin only binds to the glycon subsites, the three loops (loops 2, 3, and 4) surrounding the substrate binding pocket of MoChia1 may become more flexible when it binds to the pocket. These thermodynamic properties may help elucidate the mechanism by which allosamidin binds to different types of GH18 enzymes. This may contribute to the design of selective pesticides targeting organisms possessing GH18 enzymes.

Previous studies have shown that *M. oryzae* Secreted LyM Protein 1 (Slp1) binds to chitin oligosaccharide elicitor and suppresses the chitin-induced rice immune responses, such as those mediated by MoChia1 (27). However, in addition to binding to it, MoChia1 is capable of hydrolyzing the elicitors, which is thought to enable *M. oryzae* to evade the rice immune system more effectively. Furthermore, MoChia1 is expressed in various fungal tissues, including mycelia, spores, and appressoria, and plays a fundamental role in *M. oryzae* growth. In fact, knocking out *MoChia1* affected conidia production and germination tube development, as well as appressoria formation. In contrast, Slp1 has only been detected at the interface between the fungal cell wall and rice plasma membrane, and knocking out *slp1* did not affect appressorium formation or vegetative growth rates. These results suggest that the two proteins that bind to chitin oligosaccharide elicitors have different functions in *M. oryzae*. It is of great interest how the unique enzymatic activity of MoChia1 relates to its functions, which are not yet fully understood.

## Materials and Methods

### Materials

*N*-acetyl-D-glucosamine (GlcNAc) and chitin oligosaccharides (GlcNAc)_n_ (*n* = 2–6) were produced by acid hydrolysis of chitin (28) and purified by gel filtration on Cellufine Gcl-25 m (JNC Co., Tokyo, 3.5 × 180 cm). (GlcNAc)_2_-G, (GlcNAc)_2_-L, (GlcNAc)_2_-M, and *α*-(GlcNAc)_2_-F were prepared in accordance with previously reported methods (29–31). *N,N,N*-Triacetylchitotriitol (chitotriitol) was prepared by reduction of (GlcNAc)_3_ with sodium borohydride and purified by repeated gel filtration on Sephadex G-25. Chitin beads was obtained from New England Biolabs (Beverly, MA, USA). Ni-NTA and HiPrep 16/60 Sephacryl S-100 were the products from GE Healthcare (Tokyo, Japan). *Escherichia coli* strain 10 G cells and pRham N-His Kan vector were obtained from Lucigen (Middleton, WI, USA). *p*NP-(GlcNAc)_2_ was obtained from Megazyme (Bray, Ireland). TSKgel Amide-80 column used for (GlcNAc)*_n_* separation was from Tosoh (Tokyo, Japan). Prof. Masaru Mitsutomi, Saga University, Japan, kindly provided two mono-*N*-acetylated chitobioses, GlcN-(GlcNAc)_2_ and (GlcN)_2_-GlcNAc, which were obtained by chitinase digestion of 25–35% *N*-acetylated chitosan according to the method of Ohtakara *et al.* (32). Other reagents were of analytical grade and commercially available.

### Protein production and purification

The expression plasmid for the mature MoChia1 protein (residues 27–397 without signal sequence) with a N-terminal His_6_-tag was constructed using pRham N-His Kan (Lucigen, UK) in accordance with the manufacturer’s instructions. A synthetic gene encoding MoChia1 was obtained from Integrated DNA Technologies (IDT) (Coralville, IA, USA). The nucleotide sequence of the gene was optimized to increase expression in *E. coli* 10G without changing the amino acid sequence. Cells harboring pRham-NHis-MoChia1 were grown at 37 °C in 1 liter of Luria-Bertani medium containing 50 μg/mL kanamycin until the absorbance reached 0.8 at 600 nm. After cooling the culture to 18°C, expression was induced by adding 0.2% (w/v) rhamnose and incubation continued at 18°C for 24 hours.

### Chitinase activity

Chitinase activity of MoChia1 was measured using natural or synthetic substrates. For the *p*NP assay, 10 µL of the purified enzyme was added to 0.2 mL of 0.5 mM *p*NP-(GlcNAc)_2_ in 20 mM sodium acetate buffer (pH 5.0). After incubating the reaction mixture at 37°C for 30 min, the reaction was stopped by adding 0.8 mL of 0.25 M NaOH and the released *p*NP was quantified by measuring absorbance at 405 nm. One enzyme unit was defined as the amount of enzyme required to release 1 μmol of *p*NP per min at 37°C. For the reducing sugar assay, 10 µL of the enzyme was added to 0.25 mL of 0.2 mM (GlcNAc)*_n_* (*n* = 2–6) or 0.2 % glycol chitin in 20 mM sodium acetate buffer (pH 5.0). After incubating the reaction mixture at 37°C, the reducing power of the mixture was measured with ferricyanide/ferrocyanide reagent in accordance with the method described by Imoto and Yagishita (33). One enzyme unit was defined as the amount of enzyme producing reducing sugar corresponding to 1 µmol of GlcNAc per min at 37 °C.

### Effects of pH and temperature on the activity and stability of MoChia1

The optimal pH for chitinase activity of MoChia1 was studied by measuring the activity at various pHs (2–12) at 37°C using *p*NP-(GlcNAc)_2_ as a substrate. The buffers covering the pH scale from 2 to 12 were as follows: pH 2.0–3.0, 20 mM glycine-HCl buffer; pH 3.5–5.5, 20 mM sodium acetate buffer; pH 6.0–7.5, 20 mM sodium phosphate; pH 8.0–9.0, 20 mM Tris-HCl; pH 9.5–10.5, 20 mM glycine–NaOH, and pH 11.0–12.0, 20 mM KCl–NaOH. The effect of pH on stability of MoChia1 was studied by pre-incubating the enzyme in the buffers at various pHs at 37°C for 24 h and measuring the residual activity. The optimal temperature was determined by measuring the activity in 20 mM sodium acetate buffer (pH 5.0) at different temperatures (10 to 60 °C). The effect of temperature on the stability of MoChia1 was evaluated by pre-incubating the enzyme in 20 mM sodium acetate buffer (pH 5.0) at various temperatures for 30 min and measuring the residual activity.

### Thin layer chromatography (TLC) analysis of the reaction products produced by MoChia1

Reaction mixtures containing 2.0 μM MoChia1 and 8.0 mM (GlcNAc)*_n_* (*n* = 2–6) or its derivatives in 20 mM sodium acetate buffer (pH 5.0) were incubated at 37°C for 0.5–3.0 h. The reaction products were analyzed by TLC using TLC Silica gel 60 F_254_ (Merck, Darmstadt, Germany). The plate was developed in a solvent system comprising butan-1-ol/methanol/28.8 % ammonium hydroxide (5:4:2, by vol.) and the reaction products were detected by heating the plate after dipping it in vanillin/H_2_SO_4_ reagent.

### Anomer analysis of the reaction products produced by MoChia1

High performance liquid chromatography (HPLC) was used to analyze the anomeric forms of the substrate (GlcNAc)*_n_* (*n* = 3–6) and the hydrolysate in the reaction mixture. The enzymatic hydrolysis of (GlcNAc)*_n_* (*n* = 3–6) proceeded in 20 mM sodium acetate buffer (pH 5.0), at low temperature (25°C), to suppress mutarotation of the reaction products. The initial enzyme and substrate concentrations were 2.0 μM and 8.0 mM, respectively. After 0.5, 2, and 5 min of incubation, a portion of the reaction mixture was directly injected into a TSK Amide-80 column, which was eluted with a mobile phase comprising 70 % acetonitrile and 30 % water at a flow rate of 0.7 mL/min. The substrate and enzymatic products were detected by ultraviolet absorption at 220 nm. The cleavage site of the (GlcNAc)*_n_* (*n* = 3–6) substrate was estimated based on the distribution of the products and the anomer ratio (*β*/*α*) of the individual monosaccharide and oligosaccharide products (34,35). Since MoChia1 is a retaining enzyme, the *β*-anomer should be abundant at least at the newly generated reducing end.

### Kinetic analysis

The kinetic parameters were determined using the colorimetric assay with (GlcNAc)_2_-G at 37°C. The reaction was initiated by adding 60 µL of the enzyme in sodium acetate buffer (pH 5.0) to 60 µL of 0–20 mM (GlcNAc)_2_-G in the same buffer. The initial velocity was determined by measuring Chromogen I, which was spontaneously converted from 2-acetamido-2-deoxy-2,3-anhydro-glucopyranose (G) released from (GlcNAc)_2_-G by enzymatic hydrolysis, in accordance with the method described by Ogata et al (36). Kinetic parameters, *K*_m_ and *V*_max_, of MoChia1 were obtained by directly fitting the experimental data to the Michaelis–Menten equation using a non-linear least-fitting procedure.

### Inhibition studies

The inhibition of MoChia1 was assessed by measuring enzymatic activity with *p*NP-(GlcNAc)₂ as the substrate with allosamidin present. Reaction mixture containing 0.5 mM *p*NP-(GlcNAc)₂ and 2.5 µM MoChia1 was supplemented with allosamidin at concentrations ranging from 0 to 0.16 mM. The inhibitory concentration leading to a 50% loss of activity (*IC*_50_) was obtained by fitting the experimental data to a logistic curve using KaleidaGraph (Synergy Software).

### Isothermal titration calorimetry experiments

Isothermal titration calorimetry (ITC) experiments were performed using an iTC_200_ system (Microcal, Northampton, MA) at 30°C. Protein and buffer solutions were thoroughly degassed before use to avoid air bubbles in the calorimeter and improve results. Allosamidin solution in 20 mM sodium phosphate buffer (pH 7.0) was loaded into a syringe, and the MoChia1 solution in identical buffer, pH 7.0, was placed in the reaction cell with a volume of 0.2028 mL. For all titrations, 1 μL aliquots were injected into the sample cell with a stirring speed of 1,000 rpm. The titrations were completed after 25 injections. The integration of heat pulses and model-fitting were performed using Origin 7.0 with a single-site binding model. Individual datasets obtained from the titration experiments fitted well to the theoretical curves, providing the stoichiometries (*N*), equilibrium association constants (*K*_a_), and enthalpy changes (Δ*H*°) of the protein–ligand interactions. The binding free energy change (Δ*G*°) and entropy change (Δ*S*°) were calculated from the relationship as follows,

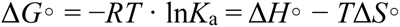

For examining the temperature dependence, ITC measurements were performed at various temperatures from 20°C to 40°C. The Δ*H*° values obtained for various temperatures were plotted against temperature, and the slope of a straight line fitted to the experimental points corresponds to the heat capacity change (Δ*C*_p_°). As the entropy of solvation is regarded as zero for proteins near 385 K, Δ*C*_p_° was converted to the solvation entropy change (Δ*S*_solv_°) at 30°C (298 K) according to the following relationship,

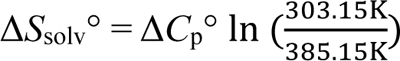

The conformational entropy change (Δ*S*_conf_°) was calculated from Δ*S*°, the solvation entropy change (Δ*S*_solv_°) obtained in this study and the mixing entropy change (Δ*S*_mix_°, − 8 cal K^−1^ mol^−1^), based on the following equation (37),

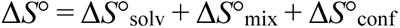

### Chitin binding assay with SDS-PAGE

The chitin-binding assay was performed using Chitin beads as the affinity matrix. Proteins (50 µg) in 20 mM sodium acetate buffer (pH 5.0) were mixed with Chitin beads (0.1%, w/v) and incubated with mixing at 25°C for 1 h. After centrifugation, the supernatant containing the unbound protein was collected and the pellet was washed with the same buffer. The chitin-bound proteins were released by boiling the affinity matrix in SDS-sample buffer and resolved on 12.5% SDS-PAGE (38). *Serratia marcescens* chitinase B (*Sm*ChiB) was prepared as described previously (39).

**Figure S1. SDS-PAGE of MoChia1.**

Lane 1: marker, Lane 2: purified MoChia1.

**Figure S2. Anomer formation catalyzed by MoChia1 from the (GlcNAc)_4_ and (GlcNAc)_6_ substrates.**

Time-dependent HPLC profiles showing the hydrolysis of (GlcNAc)_4_ (*A*) and (GlcNAc)_6_ (*B*) by MoChia1. Enzyme and substrate concentrations were 2.0 μM and 8 mM, respectively. The enzyme reaction was conducted in 20 mM sodium acetate buffer (pH 5.0) at 25°C. Numerals in the figure represent the degree of polymerization.

**Figure S3. TLC analysis of the hydrolysis of (GlcNAc)_2_-M, chitotriitol, and *α*-(GlcNAc)_2_-F by MoChia1.**

A reaction mixture comprising 2.0 μM MoChia1 and 8.0 mM of (GlcNAc)_2_-M (*A*), chitotriitol (*B*), and *α*-(GlcNAc)_2_-F (*C*) in 20 mM sodium acetate buffer (pH 5.0) was incubated for 60 min at 37°C. Reaction products were detected with methanol containing sulfuric acid. Lanes: 1, I–VI represent a standard mixture of (GlcNAc)*_n_* (*n* = 1–6); 2, substrate; 3, reaction product. The chemical structure of each substrate is shown below each TLC plate.

**Figure S4. Amino acid sequence alignment of GH18 chitinases that act on the glycosidic linkage at the reducing end of a chitin oligosaccharide.**

The alignment of MoChia1 (GenBank accession number: MGG_08054), CTBS (M95767.1) and Cfcl (XM_001400452.2) was obtained using the ClustalW (40). Dashes indicate gaps. Identical residues are indicated by the symbol "*", two identical residues out of three amino acids are marked by the symbol ":", and similar amino acids are marked by the symbol ".". The box indicates the catalytic motif (D(N)xDxE), which is conserved in GH18 enzymes. The predicted secondary structures of MoChia1 and Cfcl, as determined by Jpred and AlphaFold (41,42), are shown above and below their respective sequences. H, *α*-helix; E, *β*-sheet. The positions of the putative MoChia1 (*α*+*β*) insertion domain and the Cfcl CBM18 domain are represented by solid lines above and below the sequences, respectively.

**Figure S5. A structural comparison of CrChi-A and MoChia1.**

(*A*), CrChi-A (surface, white) in complex with allosamidin (sticks, yellow) (PDB code 4R5E). (*B*), AlphaFold model of MoChia1 (surface, cyan). The wall structure, comprised of two loops (loops 1 and 2), is surrounded by a dotted line. Allosamidin (sticks, yellow) was superimposed from the structure of CrChi-A–allosamidin complex (PDB ID 4R5E). (*C*), superimposed image of AlphaFold model of MoChia1 (cartoon, cyan) on CrChi-A–allosamidin complex. Extended loop structures (loop 1–4) found in MoChia1 were indicated. (*D*), alignment of the MoChia1 and CrChi-A sequences was obtained through PSI-BLAST analysis (43). The positions of the extended loops are indicated by solid lines above the sequence. The model of MoChia1was prepared by using AlphaFold (42). The three-dimensional structures were generated using PyMol (44).

## Data availability

No data was used for the research described in the article.

## Supporting Information

This article contains supporting information.

## Author contributions

T. O. conceptualization; T. O., T. F., and M. O. methodology; T. O., T. T., and T. F. formal analysis; S. I., C. K., T. Y., R. O., and T. T. investigation; M. O. and S. S. resources; T. O., S. I., C. K., T. Y., and R. O. data curation; T. O. and M. O. writing– original draft; T. O., T. T., T. F., S. S., and M. O. writing–review and editing; T. O., S. I., C. K., T. Y., R. O., and T. T. visualization; T. O. supervision; T. O. and M. O. project administration; T. O. funding acquisition.

## Acknowledgments

We thank the beam-line staff at BL-17A of the Photon Factory (Ibaraki, Japan) for technical assistance during data collection. This study was supported in part by a grant from the Agricultural Technology and Innovation Research Institute (ATIRI), Kindai University to T.O. In memory of Professor Masaru Mitsutomi (Saga University, Japan), who passed away during this work, we are grateful to his generous support and encouragement in this collaborative project.

## Funding and additional information

This work was supported by JSPS-KAKENHI (22K05447) and a grant from the Agricultural Technology and Innovation Research Institute (ATIRI), Kindai University to T. O.

## Abbreviations

α-(GlcNAc)_2_-F: α-(GlcNAc)_2_ fluoride
BSA: bovine serum albumin
(GlcNAc)*_n_*: *β*-1,4-linked oligosaccharide of GlcNAc with a polymerization degree of *n*
GlcNAc-GlcN: *β*-*N*-acetyl-D-glucosaminyl-(1,4)-D-glucosamine
(GlcNAc)_2_-GlcN: di-*N*-acetylated chitotriose (the reducing end residue is deacetylated)
GlcN: D-glucosamine
GlcNAc: *N*-acetyl-D-glucosamine
GH: glycoside hydrolase
(GlcN)_2_-GlcNAc: mono-*N*-acetylated chitotriose (the reducing end residue is *N*-acetylated)
(GlcNAc)_2_-G: 4-*O*-*β*-di-*N*-acetylchitobiosyl-2-acetamido-2-deoxy-2,3-anhydro-glucopyranose
(GlcNAc)_2_-L: 4-*O*-*β*-di-*N*-acetylchitobiosyl-2-acetamido-2-deoxy-2,3-anhydro-glucono-δ-lactone
(GlcNAc)_2_-M: 4-*O*-*β*-di-*N*-acetylchitobiosyl moranoline
MBP: maltose binding protein
PAMP: pathogen-associated molecular pattern
*p*NP-(GlcNAc)_2_: *p*-nitrophenyl *N*,*N*′-diacetyl-*β*-D-chitobioside
PRR: pattern-recognition receptor.

