## Supplemental Figs. for "MoChia1 is a family 18 glycoside hydrolase from the rice blast fungus, *Magnaporthe oryzae*, that releases reducing-end GlcNAc from chitin oligosaccharides"

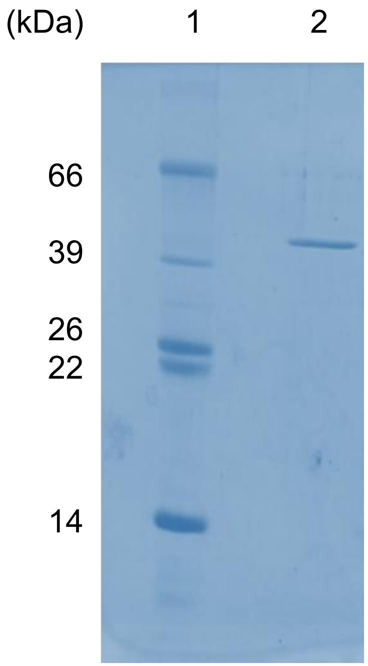


Figure S1

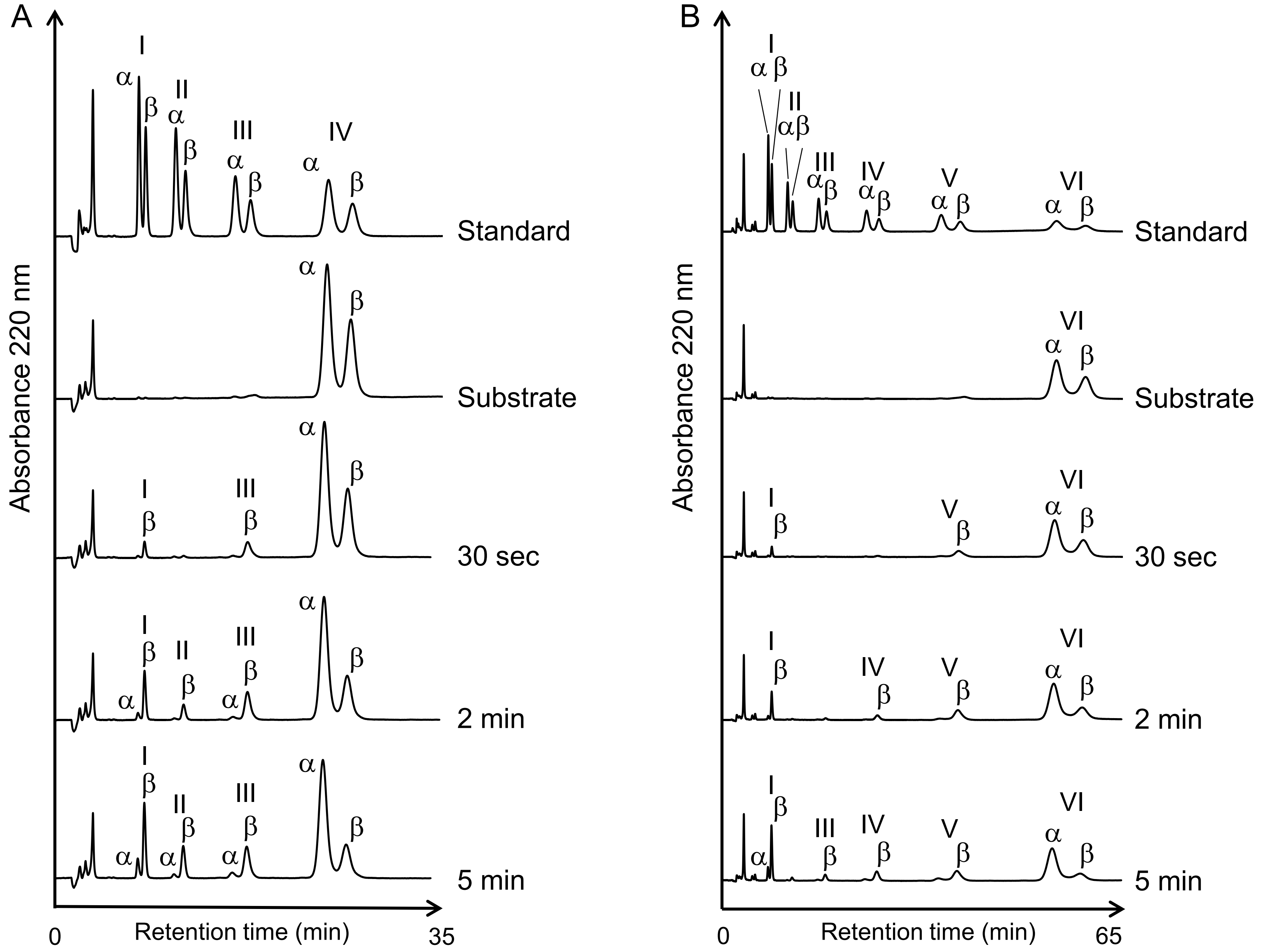
A
B
Absorbance 220 nm
Retention time (min)
0
35
65
Standard
Substrate
30 sec
2 min
5 min
I
II
III
IV
V
VI
α
β

Figure S2

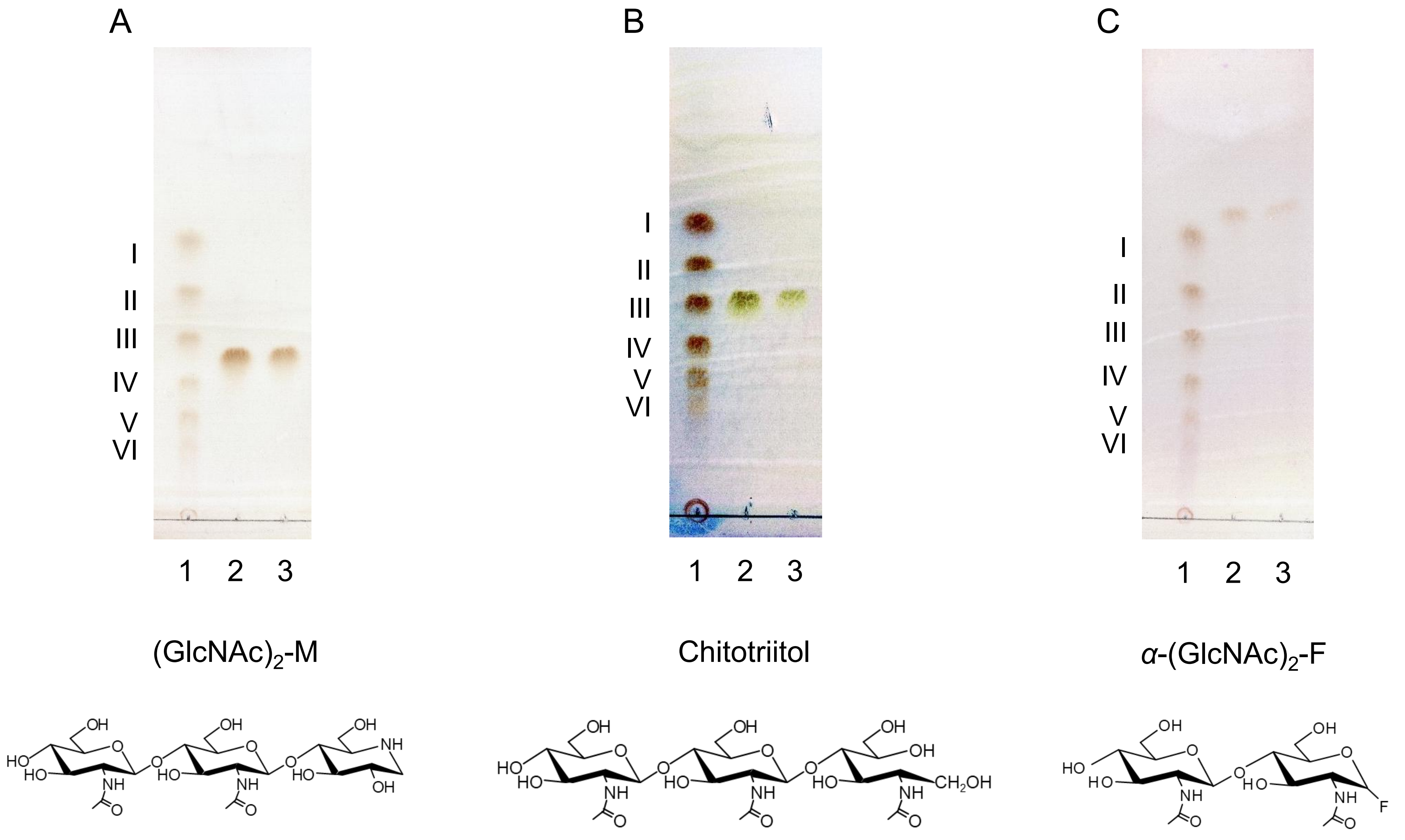
A
I
II
III
IV
V
VI
1 2 3
(GlcNAc)$_2$-M
B
I
II
III
IV
V
VI
1 2 3
Chitotriitol
C
I
II
III
IV
V
VI
1 2 3
$\alpha$-(GlcNAc)$_2$-F

Figure S3

```
                EEEEEEE                 EEEEEEEE               HHHHHHHHHH     EEEEEEE              HHHHHHHHHHHHHHHH     EEEE
MoChia1  ITPVNQSPRCVMYLTGQHN-VVPDISELRHVTHVALAFMQSPIFNEDSRKEWPLFTSVETVRRQFLPGTKVMIAIGGWGDVAGFETAAASAESRKRFAANVARMVEATGADGIDLDWEYP-119

CTBS     -----------TDCPCPEPELCRPIRHHPDFEVFVFDVGQKTWKSYDWSQITTVATFGKYDSELMCYAHSKGARVVLKGDVS--LKDIIDPAFRASWIAQKLNLAKTQYMDGINIDIEQE-107

Cfcl     -------MRFAMYVDEYHTSGLPGANQTQGITHAIMAFAQSTLFNSDSPPQFTPFESVSTMRSRFSPGTKLMIAIGGWGDTSGFSTAAKDETSRNQYAKNVAAMVNQVGFDGVDIDWEYP-113
                 EEEEEE                  EEEEEEEE              HHHHHHHHHHH    EEEEEEE              HHHHHHHHHHHHHHHH     EEEEEEEE
                          .        .   .    : . *..  . *     .     .     :  . .    :   **.:   .   .   *  :  :   :.:    **:::* *


                        HHHHHHHHHHHHHHHHHH                        HHHHHH                               HHHHHHHHH       EEEEE
MoChia1  GGNGEDYKKVPNKQREWEIEAYPLLVAEVRSAIGPDKVLSAAVPGLERDLIAFTRETVPRILRHLDFLNVMTYDLMN--RRDNVTKHHTGVQLSLDSIDAYMARGARPSQLNLGFAFY-V-236

CTBS     VN------CLSPEYDALTALVKETTDSFHREIEGSQVTFDVAWSPKNIDRRCYN---YTGIADACDFLFVMSYDEQSQIWSECIAAANAPYNQTLTGYNDYIKMSINPKKLVMGVPWYGY-208

Cfcl     GGNGQDYKTTPNSAKVSEITTYPLFLQALRSALGPNKLLSIAVPGKKEDMIAFTNTTGPQIWPSVDMVNIMSYDLMN--RRNNVTMHHTSVVDSFNTVQAYRDIGLPGNKTNLGMAYY-A-230
                        HHHHHHHHHHHHHHHHHH                                                             HHHHHHHHHH      EEEE
          .        . .       .        *.  *.:  :. * .  : *  .:.    . *    *:: :*:**  .    : ::  ::    ::   : *   .   .:  :*..:*
```

**Insertion domain**

```
           EE                                                                                                    HHHHHHHHHH
MoChia1  KYFQTR-HAECSGPGKAIGCPTLLLEDPDTGGDLGRTGGFSWHDTVP-------------------------------------------------------KDVEASFQKALRNGVYDE-300

CTBS     DYTCLNLSEDHVCTIAKVPFRGAPCSDAAGRQVPYKTIMKQINSSIS----------------------------------------------------------GNLWDKDQRAPYYN--279

Cfcl     KWFETDPSDEQECEDQPIGCKVVVMENAD-GSDNGKSGSLTFEKSVMSAPPANLKTSTDGTCGFDKGMKCPAGSCCSQYGNCGTTDDFCQAGCLSDYGTCKGISITDSWRRAQENAKLDD-305
          EEE                                                                                                   HHHHHHHHHHHEE
         .:       :       :       .:.       ::     ..::                                                              : :  .    :
```

**CBM18**

```
            EEEEE     EEEE   HHHHHHHHHHHHHH    EEEEEE        HHHHHHHHHH
MoChia1  IGGGFYYWDEEEDRWWSFDDAGAISRKFDSIFAARNLGGVFAWGLGEDAPHFEHLGAVNSRLDAITKSGAGRPETEL-377

CTBS     ------YKDPAGHFHQVWYDNPQSISLKATYIQNYRLRGIGMWNANCLDYSGDAVAKQQTEEMWEVLKPKLLQR----347

Cfcl     VMGGEYYFDDQVNVFWTWDTADMIARKFTDIVDKLGLGGVMAWSLGEDTLAWEHLGAMQEGVAAR-------------370
            EEEEEE    EEEE   HHHHHHHHHHHHHH     EEEEE        HHHHHHHHH
               * *   .    :             .    * *:  *. .      : :.  :
```

Figure S4

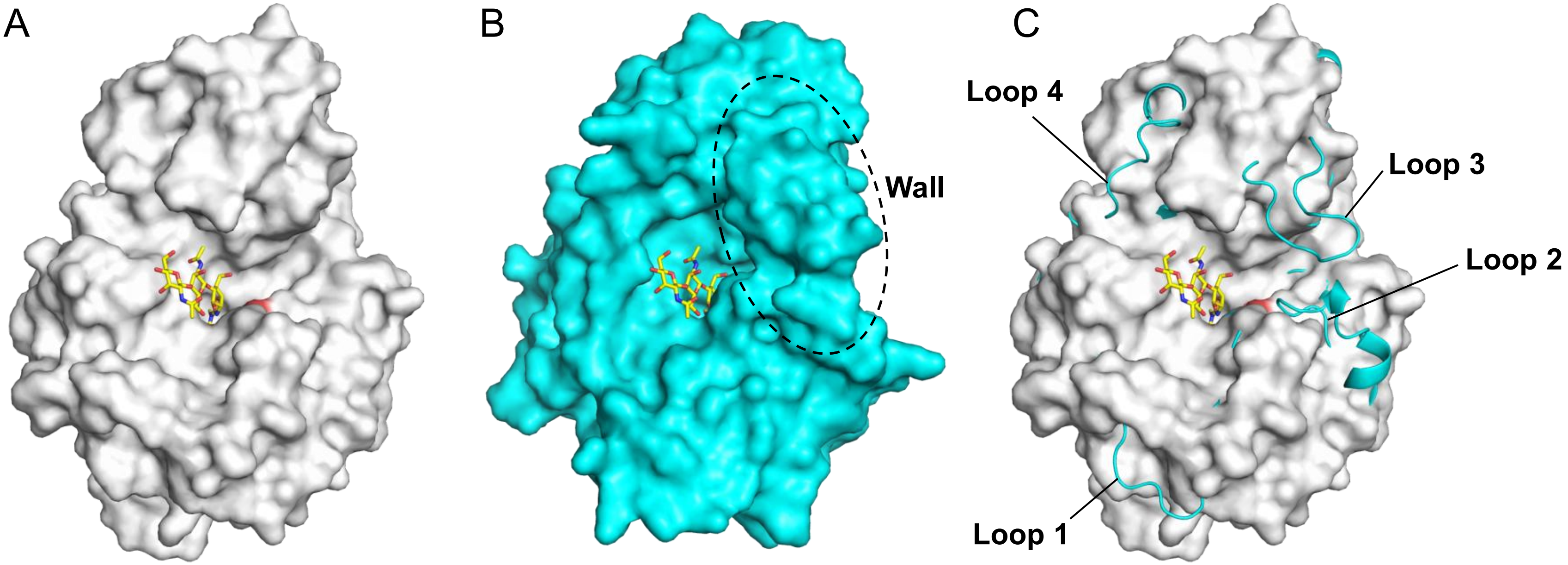


D

```
                  Loop 1
MoChia1  28   RHVTHVALAF--MQSPIFN----EDSRKEWPLFTSVETVRRQFLPGTKVMIAIGGWGDVA--GFETAAASAESRKRFAANVARMVEATGA  109
CrChi-A  26   RLFTHLYYAFAELNAPTFEVRVPPGSEKTAEDFTP--TVRR-LNPSVKTLISIGGWGSEVRDNFAKLNSDASARQRFVKSSIALARRYGF  112

                  Loop 2
MoChia1  82   DGIDLDWEYPGGNGEDYKKVPNKQREWEIEAYPLLVAEVRSAIGPDKVLSAAVPGLERDLIAFTRETVP----------RILRHLDFLNV  189
CrChi-A  83   HGLDLDYQYP-------------EPQLEMENFVKLVSELTAAIREEARTSGKPRLLLTEAVYFHQKLFPWEVVTEYPVQFIAAGLDWVNV  189

                                                                                         Loop 3
MoChia1  190  MTYDLMNRRDNVTKHHTGVQ------LSLDSIDAYMARGARPSQLNLGFAFYVKYFQTRHAECSGPGKAIGCPTLLLEDPDTGGDLGRTG  273
CrChi-A  190  MAYDFHGSWENFTGAPAALRDPNSKFTASVGIESFLAAGMPPEKLVLGIPLFGRSWLLKNNNEVG----IGAPAV--------GAGPVDG  267

                Loop 4
MoChia1  244 GFSWHDTVPKDVEASFQKALRNG---VYDEIGGGFYYWDEEEDRWWSFDDAGAISRKFDSIFAARNLGGVFAWGLGED  348
CrChi-A  250 ALSFSE---------IQNFIRGGAREVFDTTTVSAYAY--KDNVWVGYDNQQSVALKVQYAKEKR-LGGYFFWSVNQD  333
```

Figure S5
